# CTCF, BEAF-32 and CP190 are not required for the initial establishment of TADs in early *Drosophila* embryos, but have locus specific roles

**DOI:** 10.1101/2022.07.27.501678

**Authors:** Gabriel R. Cavalheiro, Charles Girardot, Rebecca R. Viales, Songjie Feng, Tim Pollex, T. B. Ngoc Cao, Perrine Lacour, Adam Rabinowitz, Eileen E.M. Furlong

## Abstract

The boundaries of Topologically-Associating Domains (TADs) are delimited by insulators and active promoters, however how they are initially established during embryogenesis remains unclear. Here, we examined this during the first hours of *Drosophila* embryogenesis. DNA-FISH on individual embryos indicates that domains form during zygotic genome activation (ZGA), but have extensive cell-to-cell heterogeneity compared to later stages. Most newly formed boundaries are occupied by combinations of CTCF, BEAF- 32 and/or CP190. Depleting each insulator from chromatin revealed that TADs can still establish during ZGA, although with lower insulation, with particular boundaries being more sensitive. Some weakened boundaries have aberrant gene expression, however the majority of mis-expressed genes have no obvious relationship to changes in domain-boundary insulation. Deletion of an active promoter (thereby blocking transcription) at one boundary had a greater impact compared to deleting the insulator-bound region itself. These results suggest cross-talk between insulators and transcription might reinforce domain formation during embryogenesis.

## INTRODUCTION

The organization of chromosomes into self-interacting chromatin domains, commonly called Topologically-Associating Domains (TADs) is a widespread feature in the animal kingdom (*1–4*). TADs are computationally defined from chromosome conformation capture data as genomic regions with enriched chromatin interaction frequencies when averaged across a population of cells (*2, 4–6*). Enhancers and their target genes are often contained within the same TAD (*3, 7*), although this organization varies from cell to cell (*5*). At some loci, the boundaries of TADs insulate regulatory elements (enhancers and promoters) within the TAD from regulatory elements in neighbouring domains, thus delimiting the space in which enhancers drive transcriptional activation (*8–11*). However, at other loci this does not appear to be the case as removal of the boundary or rearrangement of the TAD has little apparent impact on gene expression (*8, 10, 12*).

Most domain boundaries in mouse and human cells are occupied by CTCF, which binds to motifs in a convergent orientation at the two boundaries (*13, 14*). Mechanistically, TADs are formed by a loop-extrusion mechanism in vertebrates, whereby a chromatin loop is extruded by the cohesin complex until it stalls at CTCF bound regions (*15, 16*). Depletion of CTCF or components of the cohesin complex in mouse Embryonic Stem Cells (mESCs) or differentiated tissue abolishes almost all domain structure (*17–19*). Collectively, these studies demonstrate that CTCF is essential for the maintenance of the majority of TADs in vertebrates. However, it remains unclear how these domains are initially established during embryogenesis. Their formation coincides with Zygotic Genome Activation (ZGA), or just after, in Zebrafish (*20*), mice (*21*) and *Drosophila* (*22*).

In addition to CTCF, *Drosophila* has a number of other insulator proteins (*23*). Of the direct DNA binding factors, CTCF, BEAF-32, and Su(Hw) bind to the majority of domain boundaries along with the architectural cofactors CP190 or Mod(Mdg4) (*4, 24–30*). Many of these insulator proteins, including BEAF-32 (*31*) and CP190 (*32*) also bind very close to gene promoters, and a number of these factors have been proposed to function as transcriptional regulators (*31–34*). TAD boundaries also typically overlap transcribing promoters (*4, 35*), representing ∼77% of boundaries in *Drosophila* Kc167 cells (*24*), and it is currently not clear if the enrichment of insulator proteins at domain boundaries is required for boundary formation, or secondary to their role in the transcription of the boundary gene’s expression.

TADs, especially in *Drosophila*, also reflect chromatin state, which in turn reflects transcription – and partitioning of chromatin states between domains of histone acetylation and methylations (especially H3K27me3) has been proposed to lead to TAD formation in *Drosophila* (*36–38*). Actually, transcriptional state is sufficient to predict Hi-C domain structure in a number of species (*35*), including flies. However, the relative contribution of insulator protein binding, transcription or a combination of both to the formation and/or maintenance of TADs during embryonic development remains unclear.

A number of studies have begun to address this by genetic deletion or depletion of different factors *in trans*. For example, a complete deletion of CTCF (removing both the maternal and zygotic supply) *in vivo* is not required for *Drosophila* embryogenesis (*39*), or for the maintenance of TAD structure in the larval nervous system (*30*), but is required for boundary insulation and correct gene expression at a small subset of loci (*30, 39*). Depletion of BEAF-32 in Kc167 cells had little global impact on TAD structure (*24*), while depletion in BG-3 cells was reported to impact ∼20% of strong boundaries (*40*). Depletion of both CP190 and Chromator together in BG-3 cells impacted a subset of boundaries with repressive chromatin (H3K27me3), while it had little impact on active regions (*40*). Similarly, genetic deletion of CP190 impacted insulation on the subset of *Drosophila* boundaries (∼23% in Kc cells (*24*)) that do not contain active promoters, and had little impact on others (*29*).

The current lack of global regulators required for the maintenance of all TAD structure in studies that focused on late developmental stages or cell lines led us to speculate that insulator proteins are perhaps more relevant for the initial establishment of TADs in *Drosophila*, rather than for their maintenance. Here, we assessed the functional requirement for three major insulator proteins in the establishment of TADs in early *Drosophila* embryos. We first re-assessed the precise timing of domain formation in individual embryos – by measuring compaction within three TADs in single cells using high-resolution DNA-FISH. Although the genes within these loci are expressed at different stages of embryogenesis, the three TADs show similar temporal dynamics in the establishment of these domains across the very early stages of ZGA, spanning nuclear cycle (NC) 12 to 14. This confirms previous Hi-C based studies (*22, 41*), but also revealed extensive cell-to-cell heterogeneity in domain presence at this stage. We then profiled the genome-wide binding of five of the main insulator proteins (BEAF-32, CTCF, Su(Hw), CP190 and GAF) in a narrow window during the major wave of ZGA. All five proteins bind extensively along the genome, and are present in various combinations at TAD boundaries.

We depleted BEAF-32, CTCF or CP190, removing their maternal contribution, and confirmed that all three proteins are depleted from chromatin in NC14 embryos. Evaluating their requirement for the establishment of chromatin topology, using both Hi-C and DNA-FISH, revealed that the majority of TADs are still able to form in the absence of these proteins, although with lower insulation. Boundaries that are occupied by diverse combinations of insulator proteins are associated with different insulation levels at NC14, and these boundaries have different sensitivities to the loss of a single factor. Examining the impact of insulator protein depletion on gene expression revealed that ZGA occurs largely unperturbed. A few hundred genes are mis-expressed, which appears to arise through multiple mechanisms. A subset of down-regulated genes (∼1-10%) are bound by insulator proteins at their promoter, and may be regulated directly or by impacting enhancer-promoter looping, while another small subset (3-8%) are mis-regulated through enhancer-hijacking at weakened TAD boundaries. However, these are a minority – the majority (>80%) of mis- expressed genes have no obvious relationship to changes in topology and may represent more indirect effects. To disentangle the interplay between insulator proteins and transcription itself, we dissected one TAD boundary that contains an active promoter during ZGA and a bound insulator binding site. Depletion of the proteins in *trans*, and genetic deletion of the regulatory elements in *cis*, indicates that removal of the active promoter (and thereby blocking transcription) had a greater impact of TAD structure than removal of the insulator bound region itself. This suggests that transcription (or an active promoter), although not required for the establishment of TADs, may feedback and reinforce the maintenance of TAD structure as embryogenesis proceeds.

## RESULTS

### TADs are established during Genome Activation but have extensive cell-to-cell heterogeneity compared to later embryonic stages

TADs are detected by Hi-C in early *Drosophila* embryos at NC14, during the major wave of ZGA (*22, 41*). We confirmed that here, by performing Hi-C on tightly staged 2-3hr embryos (predominantly NC14) (examples in Fig. 1). To more precisely quantify the temporal dynamics and cell-to-cell variability in the establishment of TADs we performed high resolution DNA-FISH on three representative TADs with distinct features: (1) A “neutral” TAD (Fig. 1A), containing genes that are not expressed during embryogenesis, with the exception of one gene (*CG9304*), (2) the *scyl*/*chrb* TAD (Fig. 1B), containing two developmental genes (*scyl*/*chrb*) that are functionally redundant and are co-expressed during embryogenesis (*42, 43*), (3) the *tsh*/*tio* TAD (Fig. 1C), containing two essential non-redundant genes which are expressed in different patterns during embryogenesis. Both the *scyl*/*chrb* and *tsh*/*tio* TADs contain high-frequency loops, where the loop anchors are located near the developmental genes, a feature not observed in the neutral TAD. As DNA-FISH measurements are made in individual embryos and in single cells, this approach has the advantage of measuring intra-TAD proximity at absolute stages, in addition to revealing the variation in compaction across individual cells.

**Figure 1:**
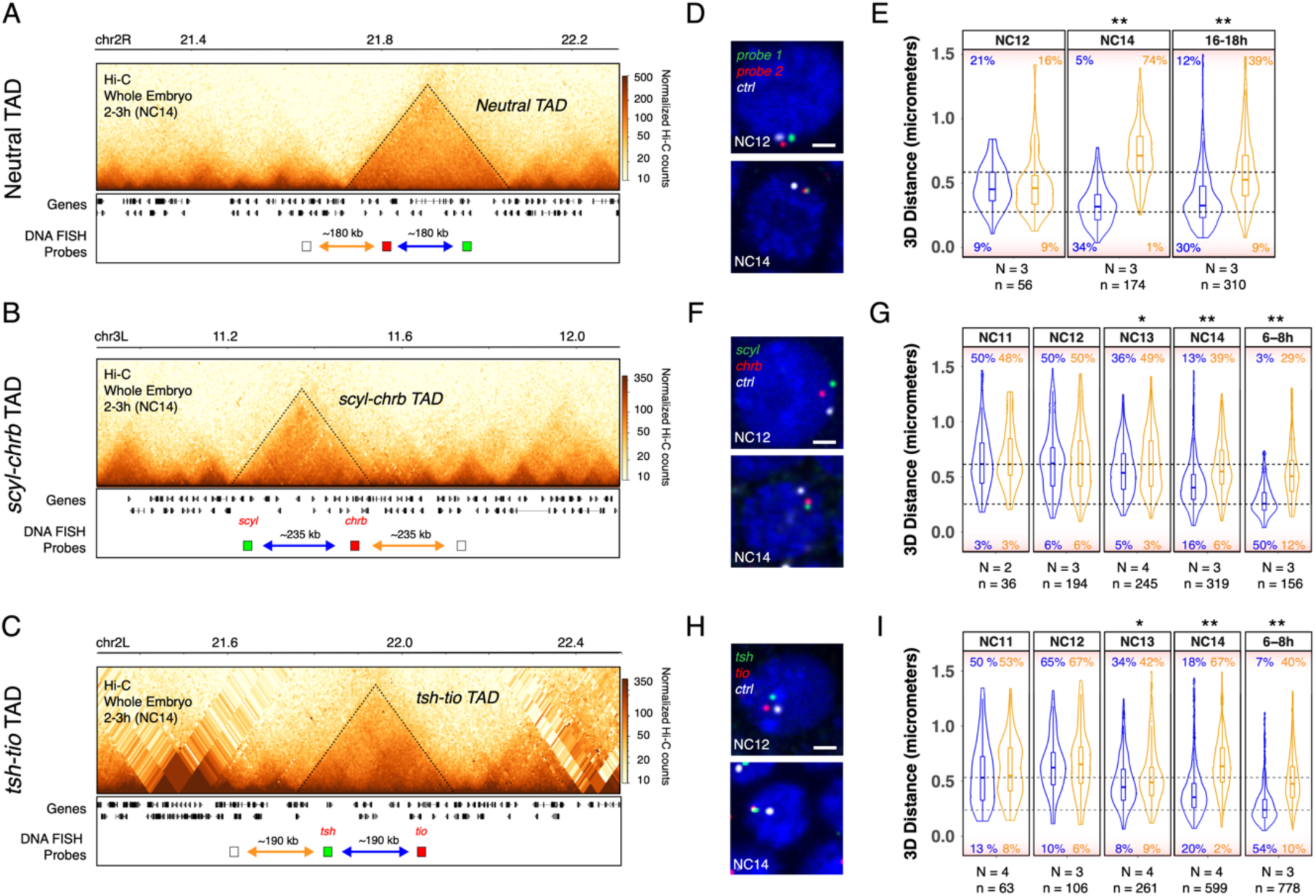
TADs are established during ZGA but have extensive cell-to-cell variation (A-C) Hi-C contact matrices from 2-3h WT embryos, showing a “neutral TAD” (**A**), *scyl-chrb* TAD (**B**) and *tsh-tio* TAD (**C**). Location of DNA-FISH probes indicated below (blue arrow: intra-TAD distance; orange arrow: inter-TAD distance). **(D,F,H)** Representative images of DNA-FISH (single confocal Z section, 100x magnification). The genomic regions targeted by FISH probes are indicated in (A-C) with the same colors (Red, green and white), with DAPI in blue. Scale bar = 1 μm. **(E,G,I)** Quantification of 3D distances between the center of mass of single spots corresponding to DNA-FISH probes in a given nucleus of embryos at each discrete stage (blue: intra-TAD distance; orange: inter-TAD distance; N = number of embryos, n = number of alleles). Dotted lines correspond to 250nm (lower) and 600nm (upper) distances between probes. Percentages indicate the number of alleles with distances >600 nm (top) or <250 nm (bottom). Kolmogorov–Smirnov (KS) test, * p < 0.01, ** p < 0.001

DNA-FISH probes spanning 7–8.5 kb were designed to target two regions within each TAD and an equidistant (in the linear genome) control probe, outside the TAD (Fig. 1A-C). For the neutral TAD, at NC12 (the minor wave of ZGA), the 3D distance between intra-TAD and inter-TAD probes is indistinguishable, with a small proportion of alleles (9%) showing overlap, defined as distances within 250 nm (Fig. 1D,E). This indicates that in the vast majority of cells (>90%) the domain is not compact, i.e. the TAD has not yet formed.

However, at NC14, the overlap between intra-TAD, but not inter-TAD, probes greatly increases, concomitantly with the major wave of ZGA: intra-TAD probes overlapped in 34% of cells compared to only 1% for inter-TAD probes (Fig. 1E). This proximity is maintained in later developmental stages (16-18h, Fig. 1E). In the *scyl/chrb* and *tsh/tio* TADs, we targeted DNA-FISH probes to the outermost anchors of the high-frequency loops, and to a control region outside of the TAD (Fig. 1B,C), and measured their 3D proximity across multiple nuclei in embryos at NC11, NC12, NC13, NC14 and 6-8h (Fig. 1F-I). At the earlier stages (NC11 and NC12), the 3D distances between the intra-TAD probes and the control probe are indistinguishable (Fig. 1G,I), similar to the neutral TAD. The compaction of the domain initiates at NC13 (as observed by the increased proximity between the intra-TAD probes), and to a much greater extent at NC14: 16% of cells vs 6% have the intra-TAD regions in close proximity (<250nm) in the *scyl-chrb* TAD and 20% vs 2% in *tsh-tio* TAD (Fig. 1G,I). In both TADs, the percentage of cells with compaction is dramatically increased as embryonic development proceeds, with 50% to 54% of cells having overlap in intra-TAD probes at 6-8h (stage 11, Fig. 1G,I).

Interestingly, the timing of TAD formation is similar across these TADs regardless of the transcriptional status of genes within the TAD. For example, the neutral TAD is formed at the same time as the two transcriptionally active TADs, supporting the hypothesis that transcription itself is globally not required for the establishment of TADs (*22*).

The single cell nature of our FISH data extends beyond the Hi-C findings by revealing extensive cell-to-cell variation in chromatin compaction (i.e. in TAD formation) among single cells across all stages (Fig 1E,G,I). To quantify this, we plotted the relative proximity between intra- and inter- TAD probe pairs, by subtracting the two (intra-TAD and inter-TAD) distances in each allele (Supplementary Fig. S1A,B,C). This revealed a considerable degree of variation among single cells in pre-ZGA embryos that decreases with time: in NC11 and NC12, the same percentage of cells display either higher intra- or inter-TAD proximity. In contrast, at NC14 most cells display higher intra-TAD proximity, indicating that TADs are forming. However, only 16-34% of cells (Fig. 1E,G,I) have intra-TAD proximity within 250nm at NC14. Therefore, although TADs are forming during the ZGA at NC14, there is still extensive cell-to-cell heterogeneity – at the *tsh/tio* TAD, for example, 18% of cells have distances >600nm at NC14, indicating that the TAD is not present in these cells at this stage (Fig. 1I, upper dashed line).

Interestingly, the cell-to-cell variability remained constant for the neutral TAD even at the end of embryogenesis (16-18hr), where 30% of cells have distances >250nm, while this changed quite dramatically for the other two TADs. In these two TADs, the number of cells with high intra-TAD proximity (<250nm) increased from 16-20% at NC14 to 50-54% at 6- 8hr of embryogenesis (Fig. 1G,I). This may reflect the transcriptional status of the genes within the two TADs (*scyl*/*chrb* and *tsh*/*tio*) which are active during these stages (in contrast to the neutral TAD), or to the presence of high-frequency structural loops, which become much more pronounced after NC14. Taken together, this indicates that cell-to-cell variation in TAD structure decreases as embryogenesis proceeds for these two TADs, while this is not the case for the neutral TAD. It also suggests that very defined TADs are not required for the regulation of gene expression during early stages of embryogenesis.

### Domain boundaries are occupied by diverse combinations of insulator proteins during the establishment of TADs

To determine how TADs are established during the major ZGA, we reasoned that insulator proteins are likely involved given the role of CTCF in TAD formation in vertebrates, the occupancy of these proteins at TAD boundaries in *Drosophila* (shown for later embryonic stages and cell lines (*24, 44, 45*)), and the availability of these proteins in early embryos due to their maternal-deposition.

We therefore first performed ChIP-seq on tightly staged NC14 embryos (2h10 – 2h40) to determine the occupancy of the four most studied *Drosophila* insulator proteins, BEAF-32, CTCF, Su(Hw) and CP190 (*23, 44*), during NC14. To our knowledge this is the first assessment of the binding of those proteins at the onset of zygotic transcription. Biological replicates for each factor are highly correlated (Supplementary Fig. S2A,B), and the DNA binding motif for each insulator protein is highly enriched under their ChIP peaks (Supplementary Fig. S2C), attesting to the quality of these NC14 ChIP datasets. All four insulator proteins have significant binding to thousands of genomic regions at NC14 (Fig. 2A,B,C): BEAF-32 - 2917, CTCF - 1319, Su(Hw) - 6134, CP190 - 5490, at < 1% IDR (Methods). Each insulator protein binds to many regions alone and in combinations with each other (Methods) (Fig. 2B,C), although interestingly the three insulators with direct DNA binding (BEAF-32, CTCF, Su(Hw)) have different distributions. BEAF-32, CTCF and CP190 combinatorial binding is more common: 51%, 58% and 60% of their peaks co- localize with at least one other insulator protein respectively, while this proportion was 34% for Su(Hw) (Fig 2C, left). The most frequent combinations are [BEAF-32 & CP190] and [Su(Hw) & CP190], representing 35% of BEAF, 20% of Su(Hw), and 41% of CP190 peaks. Triple binding is rare in comparison to single or double, and was always observed between CP190 and two other factors. Only 60 regions are co-bound by all four factors, all 60 of which are at TAD boundaries (Fig. 2C).

**Figure 2:**
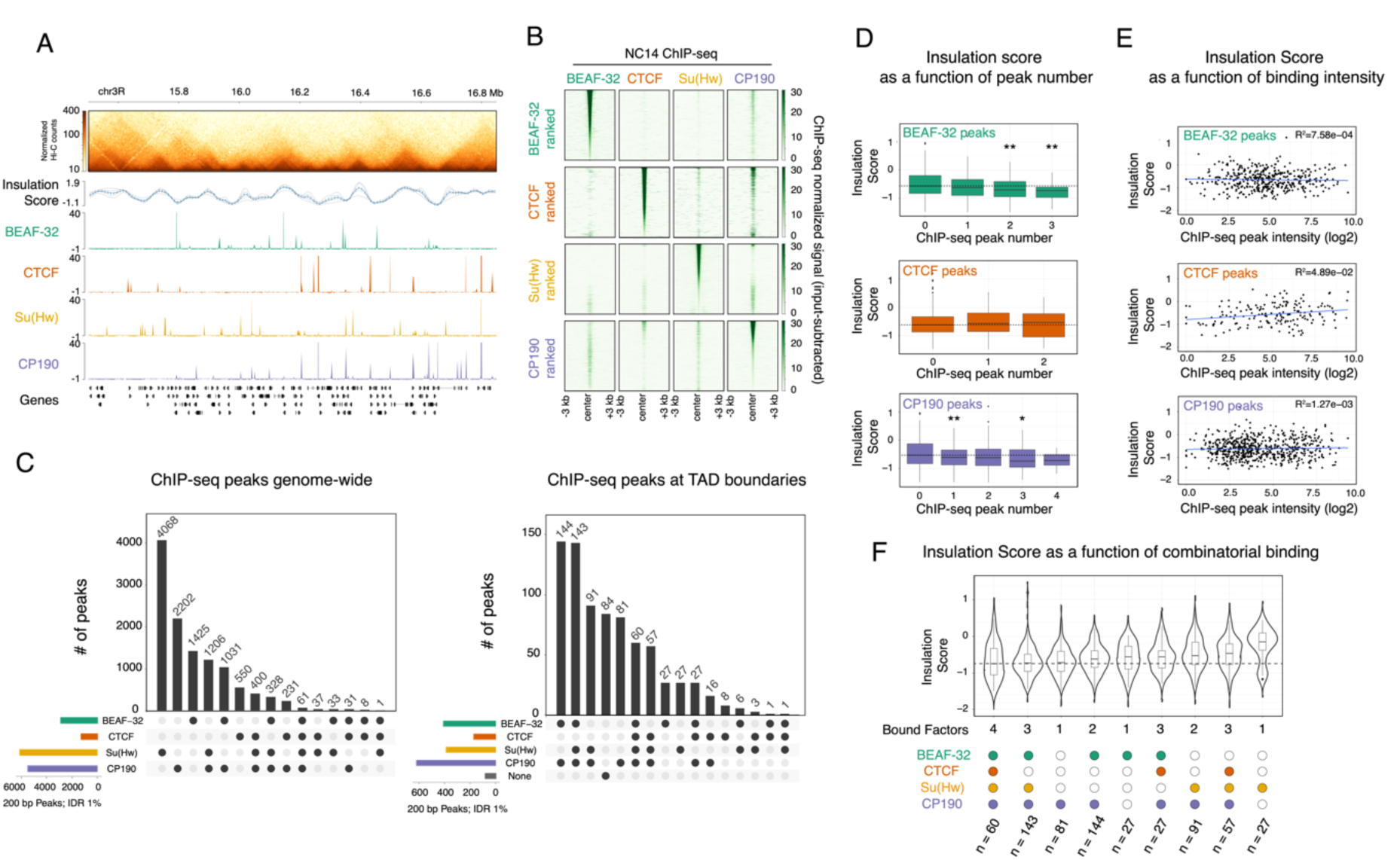
TAD boundaries are occupied by different combinations of insulator proteins during ZGA. **(A)** Hi-C matrix from 2-3h WT embryos (top), showing one genomic region with occupancy (ChIP- seq) of 4 insulator proteins at NC14. **(B)** Heatmap of insulator protein ChIP-seq normalized signal, centered at ChIP-seq peak summits. Each row is ranked by the ChIP-seq signal intensity of the indicated insulator protein, while the quantitative signal is shown for the other insulators. **(C)** UpSet plots showing the extent of insulator protein co-binding (summits within 200 bp genome-wide) (*left*) or at TAD boundaries (summits within the 10 kb boundary region) (*right*). **(D)** Box plots showing the distribution of insulation score at TAD boundaries (10 kb resolution) in 2-3h WT embryos as a function of the number of insulator ChIPs peaks at the boundary. Kolmogorov–Smirnov (KS), two- sided test * (p < 0.05), ** (p < 0.01)**. (E)** Scatter plots showing the distribution of insulation score at TAD boundaries in 2-3h WT embryos as a function of ChIP-peak intensity at the boundary. **(F)** Violin plots showing the distribution of Insulation Scores at TAD boundaries from 2-3h WT embryos as a function of insulator ChIP peaks at the boundary. Note, lower insulation score = higher insulation. Violin plots are ordered from the highest insulation (left) to lowest (right).

We also assessed the binding of the transcription factor Trithorax-like (GAF) at NC14, which was recently proposed to bind to promoter-tethering elements to establish enhancer-promoter communication in *Drosophila* (*46*). Although GAF binds to TAD boundaries at NC14 (Supplementary Fig. S2E), we observed very little co-binding between GAF and the other insulator proteins examined, including CP190 (Supplementary Fig. S2E).

CP190 is recruited to chromatin indirectly through protein-protein interactions with other insulator proteins and therefore does not have a canonical DNA binding motif (*47–49*). This is reflected in the extensive co-localisation of CP190 with the other three insulator proteins (Fig. 2C, 60% of peaks overlap). This is in keeping with the *de novo* motif enrichments under CP190 peaks, where Su(Hw), CTCF and BEAF-32 motifs are within the first five most enriched motifs (Supplementary Fig. S2C). Motif analyses of the 40% of CP190-only peaks identified additional putative recruiters of CP190 during these early stages of embryogenesis (Supplementary Fig. S2D), including the insulator protein Pita (*50*), and the transcriptional regulators Knirps, Jim, Nautilus and Visual system homeobox 2.

We next assessed how insulator occupancy relates to the newly established TAD boundaries during ZGA. As the ability to define TAD boundaries in a given condition varies greatly depending on the resolution of the Hi-C matrices, and the TAD calling algorithm used, we used our Hi-C data from tightly staged 2-3hr embryos to call TADs at multiple base-pair resolutions (2, 5, 10 kb) and q-values (0.1, 0.05 and 0.01) (Supplementary Fig. S3A). The 10 Kb resolution and q-value 0.1 gave the most consistent results, showing a better visual overlap with TADs and avoiding splitting larger domains. By using these thresholds we identified 772 high confidence TAD boundaries in 2-3hr embryos (Supplementary Fig. S3A,B, Fig. 2A). Approximately 90% (703/772) of these domain boundaries have at least one of the four analyzed insulator proteins binding within the 10kb boundary window at NC14 (Fig. 2C, right), with CP190 being the most frequent (80%, 624/772). TAD boundaries are preferentially bound by combinations of insulators rather than single proteins, with 70% of all boundaries being occupied by two or more of the four insulators (Fig. 2C, right). This proportion is likely an underestimate given that there are other *Drosophila* insulator proteins reported that we did not profile here. The most prevalent combinations are [BEAF-32 & CP190] and [BEAF-32, Su(Hw) & CP190], present in 17% and 18% of TAD boundaries, respectively (Fig. 2C, right).

Two trends were previously reported in mice or in *Drosophila* cell lines. In mammals, both a higher number and higher intensity of CTCF ChIP peaks was proposed to provide robustness to TAD boundaries (*12, 51*). Here, during the establishment of TADs in early *Drosophila* embryos we observed a similar trend for stronger insulation at boundaries with multiple BEAF-32 or CP190 peaks, although this does not hold true for multiple CTCF peaks (Fig. 2D, note a lower insulation score reflects higher boundary insulation). Moreover, ChIP peak intensity at TAD boundaries for either BEAF-32, CTCF or CP190, is not correlated with stronger insulation (Fig. 2E). In *Drosophila* Kc cells, the strength of TAD boundaries (as measured by insulation score) is correlated with the number of co-bound insulator proteins (*25*). We observe a similar trend in NC14 embryos: 26% (202/772) of domain boundaries are occupied by either all four (60) or by three (BEAF-32, Su(Hw) & CP190) factors (Fig. 2C, right), and these are among those with the highest insulation (Fig. 2F). For boundaries occupied by other combinations of three or less insulator proteins, the insulation score varies depending on the identity of the bound insulators (Fig. 2F). For example, boundaries occupied by [BEAF-32, CTCF & CP190] or by [CTCF, Su(Hw) & CP190] have weaker insulation than those bound by [BEAF-32, Su(Hw) & CP190] (Fig. 2F). Similarly, insulation strength varies in boundaries occupied by different combinations of two insulator proteins depending on their identity: for example, the median insulation of [BEAF-32 & CP190] occupied boundaries is between the medians of the triple combinations [BEAF-32, CTCF & CP190] and [BEAF-32, Su(Hw) & CP190]. CP190-only boundaries are an interesting exception, which have almost as strong insulation as those bound by all four proteins (Fig. 2F), suggesting that other CP190-recruiters are important for insulation at this stage of embryogenesis.

In summary, our results indicate a strong diversity in the occupancy of TAD boundaries by combinations of insulator proteins at NC14. The binding of more proteins has a higher likelihood that a site will form a boundary. However, it is not simply the number of bound insulator proteins that is the sole determinant of insulation; the nature of the bound factors (and presumably the genomic context) influences the strength of insulation at a boundary.

### Genetic depletion leads to very efficient removal of insulator proteins from chromatin in early embryos

To investigate the functional contribution of each insulator protein to the establishment of TADs and gene expression in early embryos, we removed the maternal deposition of each protein in females in the developing oocyte. For CTCF, we used our previously generated knockout allele that completely removes both the maternal and zygotic CTCF mRNA and protein (*39*). Genetic knockout of BEAF-32, Su(Hw) or CP190 leads to strongly reduced fertility, sterility or homozygous lethality, respectively (*52–54*). Thus for these factors we used an alternative depletion strategy, through RNAi-mediated knockdown in the female germline (*55*). We could successfully obtain embryos after knock down of maternal BEAF- 32 or CP190 in the female germline. Unfortunately, knock down of maternal Su(Hw) led to female sterility, and we could therefore not obtain embryos to study the contribution of Su(Hw) in the establishment of TADs. We therefore focused on the role of CTCF (using genetic deletion of the maternal and zygotic contribution) and BEAF-32 and CP190 (by RNAi knockdown).

The efficiency of protein depletion in NC14 embryos was assessed by two metrics. First, by western blot, which indicates that the three insulator proteins (BEAF-32, CTCF or CP190) were very strongly depleted to almost undetectable levels (Fig. 3A,B). Second, by quantitative CUT&Tag (C&T) using spike ins, to determine the protein’s depletion from chromatin. For both wild-type (WT) and insulator-depleted embryos, C&T was performed on 50,000 *Drosophila melanogaster* nuclei (isolated from NC14 embryos) combined with 50,000 nuclei from another *Drosophila* species (*D. virilis*) (isolated from 2-4 hr embryos).

**Figure 3:**
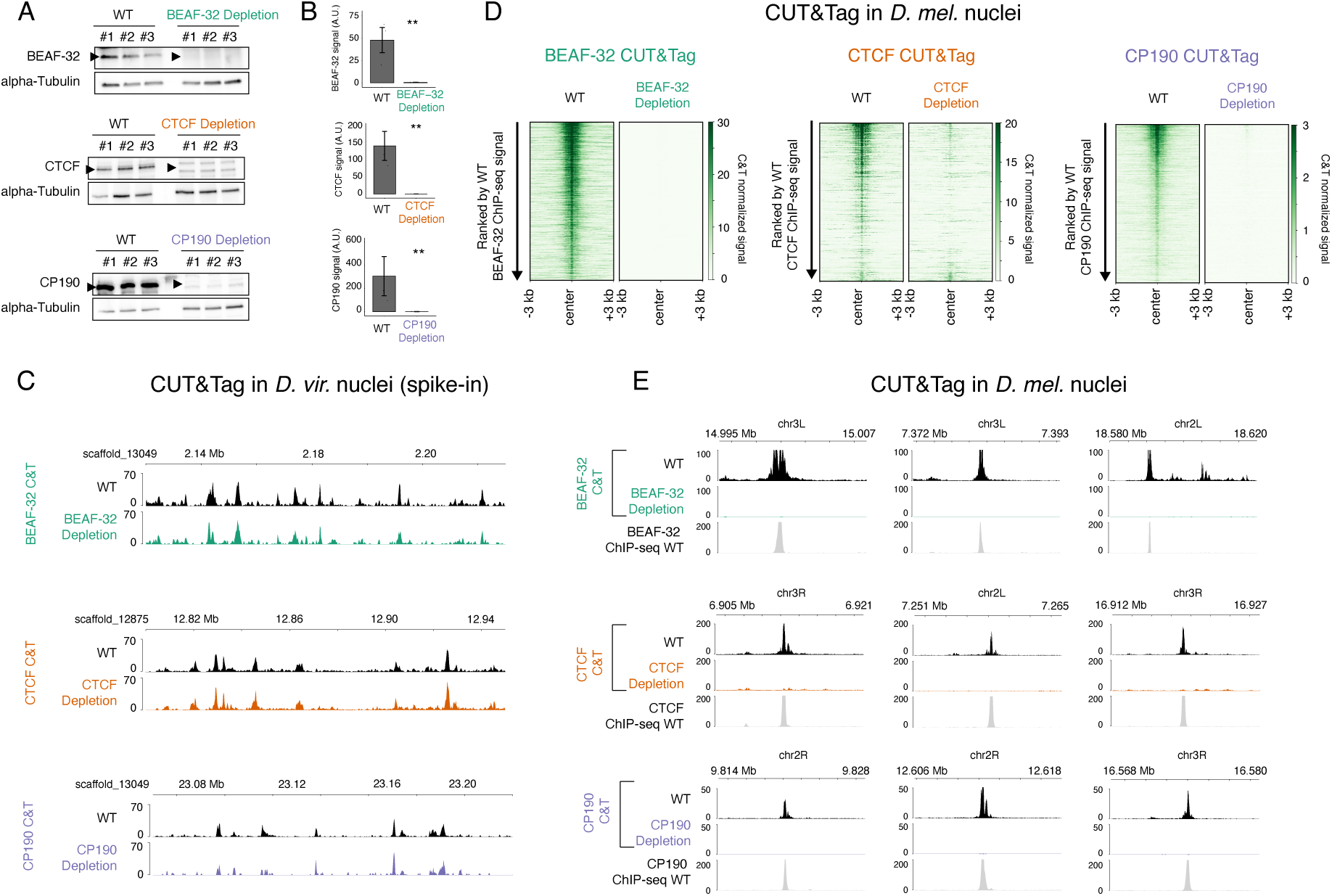
Efficient depletion of insulator proteins from chromatin. **(A)** Western blots for BEAF-32 (top), CTCF (middle) and CP190 (bottom) comparing three biological replicates of NC14 WT vs. insulator depleted embryos after maternal depletion of each insulator protein. BEAF-32, CTCF and CP190 bands indicated by arrow. Alpha-Tubulin was used as a loading control to normalize all experiments. **(B)** Quantification of the Western blots, showing almost complete depletion. Two tailed T-test, ** p < 0.001. **(C)** Genome browser showing *D. virilis* CUT&Tag (C&T) signal at representative regions from experiments where nuclei were pooled (spiked-in) with *D. melanogaster* WT and insulator-depleted nuclei. In both samples, the *D. virilis* insulator peaks are present, confirming that the C&T worked. **(D)** Heatmaps of C&T normalized signal (with *D. virilis* rescaling) in *D. melanogaster* nuclei of WT and insulator-depleted samples from NC14 embryos, centered on the ChIP-seq peak summits for each protein. Heatmaps are ranked by C&T peak signal in WT embryos. (*left*) BEAF-32 C&T, (*middle*) CTCF C&T, (*right*) CP190 C&T. **(E)** C&T signal (after *D. virilis* rescaling) in *D. melanogaster* showing representative regions in WT and insulator-depleted embryos. ChIP-seq in WT embryos, showing peaks co-localized to C&T signal in WT embryos. These representative loci show the clear absence of insulator chromatin binding in depleted embryos compared to WT.

Importantly, this *D. virilis* spike in was added to every sample (both WT and insulator- depleted *D. melanogaster* nuclei) to control for the efficiency of tagmentation between samples and thereby enables a more accurate quantification of the reduction in ChIP peaks. Such spike-in controls are particularly important in cases like this where we expect almost a complete loss of binding in the depletion condition.

The occupancy profiles from the spiked-in *D. virilis* nuclei for all three insulator proteins were nearly identical for nuclei pooled with *D. melanogaster* WT and insulator-depletion matched samples, indicating that all C&T experiments worked efficiently (Fig. 3C). In contrast, in *D. melanogaster* nuclei there was a dramatic reduction in chromatin binding for BEAF-32, CTCF and CP190 in their respective depletion conditions (Fig. 3D,E), while the binding was still present in the pooled *D. virilis* nuclei (Fig. 3C). We could detect nearly no significant peaks overlapping any of the three insulator proteins’ WT peaks following their depletion (Fig. 3E). This is important, as even low levels of insulator protein is enough to sustain chromatin topology in other models (*17, 56*). We did observed some non-specific peaks in the insulator depleted samples not present in WT conditions. As these were only present after the depletion of the protein (i.e. in the *D. melanogaster* nuclei) and not in the *D. virilis* nuclei, this is likely spurious Tn5 activity (ATAC-like signal) in the absence of the correct epitope for the primary antibodies. However, this is purely a technical artifact, while importantly the binding of BEAF-32, CTCF and CP190 at endogenous wild-type peaks is severely reduced or completely absent (Fig. 3D,E).

These results confirm that our depletion conditions are very efficient at removing these factors from their bound regions and therefore can be used to assess the function of BEAF-32, CTCF and CP190 in the establishment of TADs.

### Insulator depletion does not inhibit TAD formation, but does weaken specific boundaries with different properties

To assess the requirement of these insulator proteins in the establishment of TADs, we performed Hi-C in embryos maternally depleted of each insulator at NC14. Surprisingly, depletion of BEAF-32, CTCF or CP190 did not prevent the establishment of the majority of TADs, as observed in the Hi-C matrices (Fig. 4A). We detected a similar number of TADs across all genotypes (Fig. 4B), which is robust to different TAD calling parameters (Supplementary Fig. S3B (*24*)). Chromosomal interactions across genomic distances are also similar for all genotypes, both wild-type and the different insulator-depleted embryos (Supplementary Fig. S3C). These results highlight the overall stability of topology at the TAD-level and at very long range compartmental-level interactions. However, quantifying the insulation scores of TAD boundaries revealed a subtle genome-wide decrease in insulation (i.e. an increase in insulation score) in all three depletions (Fig. 4C), even though most boundaries are still detectable. Similarly, quantifying interaction frequencies within TADs and across the boundary with the neighbouring TADs (intra- vs inter-TAD interactions) revealed that all three insulator depletions have higher inter-TAD interactions compared to WT (Supplementary Fig. S3D). Although significant, this difference between the depletions and wildtype is subtle. Furthermore, all three depletions are more highly correlated between each other compared to the WT replicates (Supplementary Fig. S3E).

**Figure 4:**
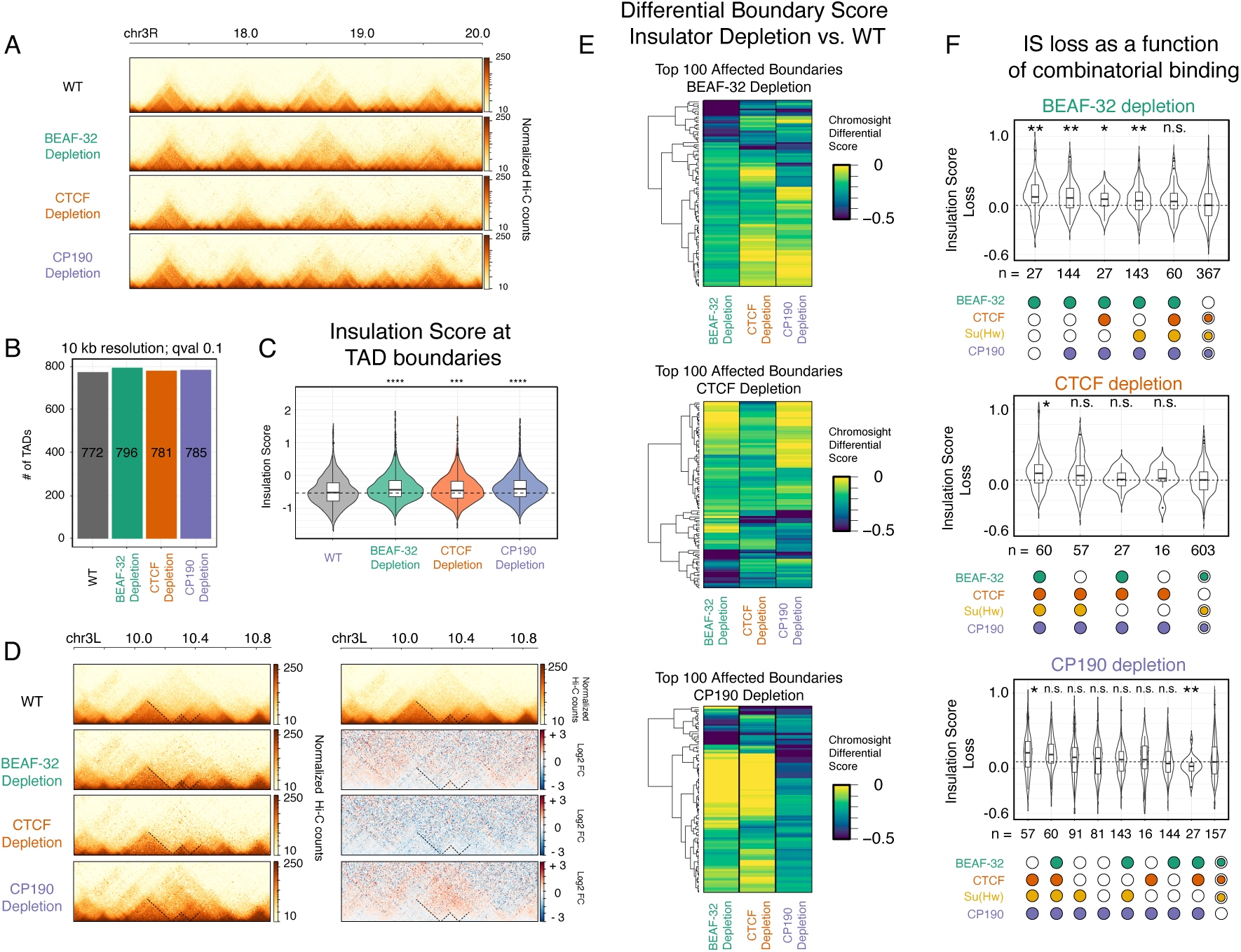
Depletion of insulator proteins has little global impact on the establishment of TADs during ZGA. **(A)** Hi-C matrices from WT and insulator-depleted embryos (2-3h), normalized by read counts, showing a representative region on chr3R. **(B)** Quantification of total TAD numbers called in WT and insulator-depleted samples, using Hi-C explorer (*24*) with matrices at 10 kb resolution. **(C)** Quantification of insulation score at all TAD boundaries in WT vs. insulator depleted embryos. Kolmogorov–Smirnov (KS) test, **** p < 0.0001. **(D)** *Left*: Hi-C matrices of one of the top-100 regions with a weakened TAD boundary in CP190 depleted embryos. *Right*: differential Hi-C signal (log2 Fold-Change) of the same region, where the Hi-C signal (interaction frequencies) from the insulator depleted samples is divided by the WT signal. Red = higher signal in depleted samples, blue = higher signal in WT embryos. **(E)** Heatmaps of the differential scores (chromosight) (*57*) of observed changes at TAD boundaries in insulator depletion vs. WT embryos, for the top 100 weakened boundaries per genotype. Negative values (blue) indicate weakened boundary in comparison to WT, while zero (yellow) indicates no change. **(F)** Distribution of insulation score (IS) loss at TAD boundaries in BEAF-32, CTCF and CP190 depleted embryos (upper, middle and lower panels) compared to WT embryos at 2-3hr. Plots ordered by highest loss of insulation, left, to smallest, right. Boundaries occupancy by insulator proteins at NC14 (in WT embryos) shown underneath. Boundaries not occupied by the depleted insulator shown on the right. Kolmogorov– Smirnov (KS), two-sided test. N.s. (p > 0.05), * (p < 0.05), ** (p < 0.01)

In addition to this genome-wide trend for lower domain boundary insulation, some individual loci appear more strongly affected. For example, some loci have weekend boundaries in specific genotypes, which are observed as gains of interactions across the TAD boundary, as seen in the differential Hi-C matrices (Fig. 4D). To systematically identify regions exhibiting differential interactions (disrupted TAD boundaries), we used a computer-vision based algorithm, Chromosight (*57*), which identified 148/878 (17%), 119/869 (14%) and 151/878 (17%) boundaries with differential interactions (using a cutoff of < - 0.1) between WT and BEAF-32, CTCF or CP190 depleted embryos, respectively (Table S1). This changed to 43 (4.9%), 43 (4.9%) and 81 (9%) of boundaries using a more stringent cut off <-0.2 (Table S1). Clustering the top 100 disrupted boundaries in each genotype shows the highest reduction in differential boundary score (from chromosight), for that proteins depletion, as expected (Fig. 4E, less yellow). This also revealed a number of boundaries whose insulation is affected by more than one genotype (Fig. 4E, green/blue). We complemented this with a visual curation of all affected TAD boundaries. A large fraction of the computationally identified top 100 affected TAD boundaries have an obvious alteration in the contact matrix by visual inspection (60, 52 and up to 75 for BEAF-32, CTCF and CP190 depletions, respectively; Table S1). CP190 depletion had the strongest effects on boundary loss (as also seen in Fig. 4D, and subsequent figures), consistent with its presence at ∼80% of all boundaries (Fig. 2), and its proposed insulator cofactor role.

To determine if there are specific features of TAD boundaries that are sensitive to different insulator protein depletion, we divided boundaries into different classes based on their combinatorial insulator binding (as in Fig. 2) and analyzed which classes had the largest loss in insulation score. The violin plots are ordered from the largest loss of insulation (greatest change in insulation score, Fig 4F, left) to boundaries that are not bound by that insulator protein for comparison (Fig. 4F, right). This revealed that depletion of BEAF-32, CTCF or CP190 led to a stronger reduction in insulation at domain boundaries occupied by these respective factors (Fig. 4F, note higher insulation score loss indicates a larger decrease in boundary insulation), which although expected confirms the specificity of the depletions. It also revealed interesting differences between BEAF-32 and CTCF & CP190. The boundaries that are only bound by BEAF-32 (or [BEAF-32 & CP190]) are the most severely affected after BEAF-32 depletion compared to boundaries co-occupied by other factors or to boundaries not occupied by BEAF-32 (Fig. 4F). Conversely, depletion of CTCF or CP190 has more effect on combinatorially bound boundaries. For example, depletion of CTCF had a stronger reduction in insulation at boundaries bound by all four insulator proteins, while CP190 had a stronger impact on boundaries occupied by [CTCF, Su(Hw) & CP190], suggesting that both insulators function by more cooperative interactions (Fig. 4F).

These differences between BEAF-32, CTCF and CP190 are also apparent when looking at the relationship between the changes in insulation score and the number of ChIP peaks for that factor. TAD boundaries that overlap more BEAF-32 peaks (within the 10kb window) in WT embryos are more dependent on that insulator protein (i.e. had a larger reduction in insulation after depletion) (Supplementary Fig. 4A). However, this trend is not significant for CTCF and CP190 (Supplementary Fig. 4A), and there is no correlation between the ChIP peak height and insulator score for all three proteins (Supplementary Fig. 4B).

Examining the distance of an affected boundary to the closest insulator ChIP peak also revealed differences between each insulator protein. Boundaries affected in CTCF and CP190 are closer to a CTCF peak, and further away from a BEAF-32 or CP190 peak in comparison to a random background of unaffected boundaries (Supplementary Fig. 4C). In BEAF-32 depletion, affected boundaries are further away from CP190 peaks than random, but not significantly closer to BEAF-32 peaks. Boundaries affected in all three depletions were further from CP190 peaks (Supplementary Fig. 4C), which might be due to the widespread presence of CP190 in 80% of boundaries (and thus present in many non-affected boundaries). In most cases, the significant difference in the distance distributions concentrates at about 50Kb from the TAD boundaries, and is therefore difficult to reconcile with a simple direct relationship between loss of binding at the boundary and loss of boundary insulation, but again hints at a functional connection between CTCF & CP190.

Taken together, these results indicate that the initial establishment of TADs during *Drosophila* embryogenesis is generally robust to the loss of a single insulator protein. However, TAD boundary insulation is globally decreased, and some specific TAD boundaries are more sensitive to the loss of a given insulator protein than others. Our observations indicate that CP190 and CTCF function more combinatorially compared to BEAF-32, even though BEAF-32 and CP190 co-bind much more extensively throughout the genome.

### Depletion of insulator proteins leads to transcriptional defects through different mechanisms

To investigate how insulator protein depletion during the establishment of TADs affects gene expression, we performed RNA-seq in manually-selected embryos at NC14, in triplicates. Principal Component (PC) analysis illustrates that in all three genotypes, PC1 clearly separates depletion and WT replicates (Supplementary Fig. S5A). To avoid any potential confounding *trans*-effects, we removed maternally-deposited genes from the analysis (using RNA-seq data from unfertilized oocytes (*10*)) and thereby focused only on genes that start to be expressed at the ZGA. Examining these strictly zygotically expressed genes identified 325, 436 and 597 Differentially Expressed Genes (DEGs) in BEAF-32, CTCF and CP190 depleted embryos, respectively compared to staged matched wild-type embryos (| log2 Fold Change (FC) | > 0.7 & FDR < 0.05) (Supplementary Fig. S5B, Table S2). This is largely balanced between the number of up and down regulated genes (Supplementary Fig. S5B), and they are not enriched in any particular biological function (looking for GO-term enrichments). Around 25% of all DEGs (263/1030) have expression changes in the same direction in two of the three genotypes. Thus, the transcriptional responses following depletion of insulator proteins are largely genotype-specific. However, CTCF and CP190 depletion resulted in slightly more overlap in mis-expressed genes (Fig. 5A), in keeping with the similarity in their most affected TAD boundaries (discussed above (Fig. 4)).

**Figure 5:**
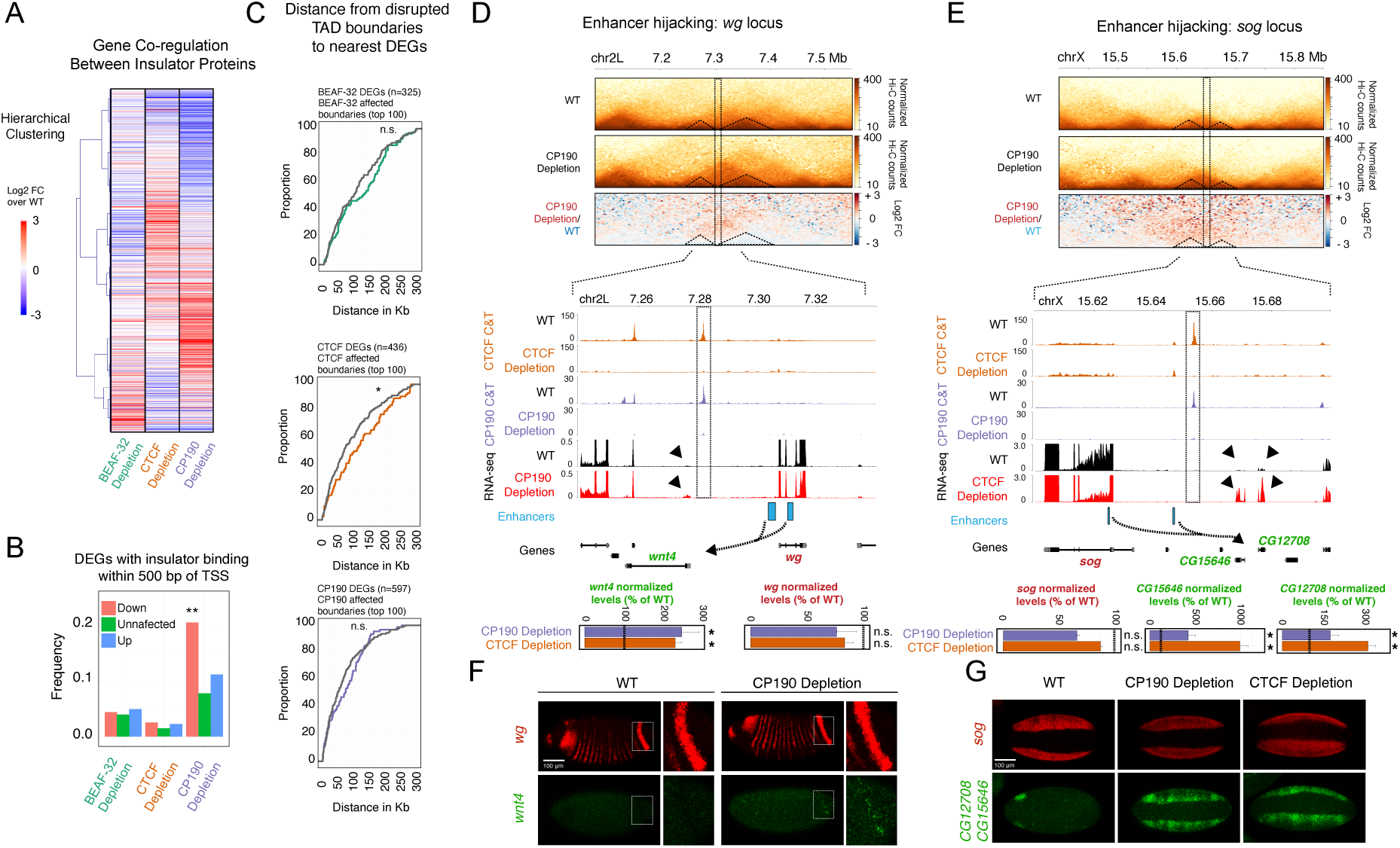
Transcriptional changes after insulator protein depletion. **(A)** Heatmap of RNA-seq changes (log2FC) in insulator depleted vs. WT NC14 embryos, with increased (red) and decreased (blue) changes shown, clustered by hierarchical clustering. Only zygotic differentially expressed genes (DEGs) in at least one genotype are included – 1030 genes. **(B)** Quantification of the proportion of DEGs in which the promoter region (+/- 500 bp from the TSS) overlaps a ChIP-seq peak for the corresponding insulator protein (in WT embryos). Different colors represent DEGs (up- and down-regulated) or unaffected genes. Fisher’s exact test ** p < 0.0001. **(C)** Cumulative curves of the distance between the promoter of DEGs after each insulator depletion (colored lines) to the nearest disrupted TAD boundary. Distances for random non-affected boundaries shown in grey. Kolmogorov–Smirnov (KS), two-sided test, n.s. (p > 0.05), * (p < 0.05). **(D, E)** Two loci showing enhancer hijacking: *Top*: Hi-C matrices and differential Hi-C matrices (log2 FC) of the *wg* (**D**) and *sog* (**E**) loci. Increased interactions after insulator depletion (red) vs. WT (blue). *Bottom*: Zoom in of CP190 and CTCF occupancy (C&T) in WT and insulator-depleted NC14 embryos and RNA-seq signal from WT (black) and insulator-depleted (red) embryos. Experimentally characterized enhancers are shown in blue, TADs separated by a weakened boundary highlighted with dotted triangles. Potential new regulatory connections (“enhancer hijacking”) represented by dotted arrows. Genes highlighted in green are upregulated in either CP190 (*wnt4*, *CG15646*) or CTCF (*wnt4*, *CG15646*, *CG12708*) depletion presumably due to enhancer hijacking, while the expression of the enhancer’s original target genes (highlighted in red) are not affected. RNA-seq normalized levels of the highlighted genes in CTCF and CP190 depleted embryos are displayed in a bar plot; non-significant (n.s., FDR > 0.05), * (FDR < 0.05). **(F)** *In-situ* hybridization of *wg* (red) and *wnt4* (green) in WT (left) and CP190-depleted (right) NC14 embryos. Inset highlights the posterior stripe of *wg* expression, where a few cells in CP190-depleted embryos ectopically express *wnt4*. The colors match the gene names in (C). Scale bar = 100 μm. **(G)** *In-situ* hybridization of *sog* (red) and *CG12708/CG15646* (green) in WT (left), CP190-depleted (center) and CTCF-deleted (right) NC14 embryos. Colors match the gene highlighting in (D). *CG12708/CG15646* are ectopically expressed in a similar pattern as *sog*, suggesting mis-regulation by *sog* enhancer(s). Scale bar = 100 μm.

We also examined the expression of genes that are activated during the minor and major waves of Zygotic Genome Activation (*58*), and are involved in the first spatial patterning events of the embryo. The majority of these genes had no significant changes in their mRNA levels after the depletion of BEAF-32, CTCF or CP190, with a few exceptions including the upregulation of the homeotic gene *scr* in CTCF mutant embryos (Table S3). This indicates that ZGA and the activation of the majority of the anterior-posterior and dorsal-ventral patterning genes (some of which are regulated by known distant-acting elements e.g. (*59–61*)) does not require these insulator proteins.

Insulator proteins have also been implicated in directly regulating gene expression (*62, 63*). In *Drosophila*, for example, BEAF-32 (*31*) and CP190 (*32*) were proposed to have an activator role by directly binding to a subset of promoters, while CTCF was proposed to have either a transcriptional repressor (*64*) or activator (*17, 63, 65*) function. A subset of DEGs have insulator protein binding directly at their promoter (+/- 500bp from TSS) at this stage of embryogenesis (ZGA NC14 embryos) (Fig. 5B): 5% (15/325) of BEAF-32, 2% (10/436) of CTCF and 16% (94/597) of CP190 DEGs. For BEAF and CTCF, these DEG were roughly balanced between genes that were up or down-regulated (7/15, 4/10 respectively). However, for CP190 65% (61/94) of these genes with binding at the promoter have reduced expression upon depletion (Fig. 5B). Therefore, our results show little evidence for direct promoter regulation by BEAF-32 or CTCF at these stages, however CP190 may regulate the expression of a subset of genes more directly, independently of a role in TAD formation (Fig. 5B). This could involve the regulation of promoter-enhancer loops, or a more direct regulatory role on Pol II activity at the promoter. However, this represents only a small fraction of all mis-expressed genes after CP190 depletion.

To explore the relationship between TAD boundary disruption and gene misexpression, we focused on the top 100 most affected TAD boundaries in each genotype, and measured their distance to the nearest DEG (Fig. 5C). In all three depletions, the nearest DEG is located up to 300 kb away from a disrupted boundary, with a median distance of ∼125, 100 and 70 kb for BEAF-32, CTCF and CP190, respectively (Fig. 5C). This distance distribution is comparable to the background (i.e. the distance of 100 unaffected boundaries to the nearest DEG) (Fig. 6C, compare coloured line (disrupted TADs) to grey (background)), and is larger than the average TAD size that most of these genes reside in. This indicates that DEGs are generally not enriched near disrupted TAD boundaries, and that the TAD boundary is not constraining enhancer activity in the majority of cases, as we and others observed previously (*8, 10*).

**Figure 6:**
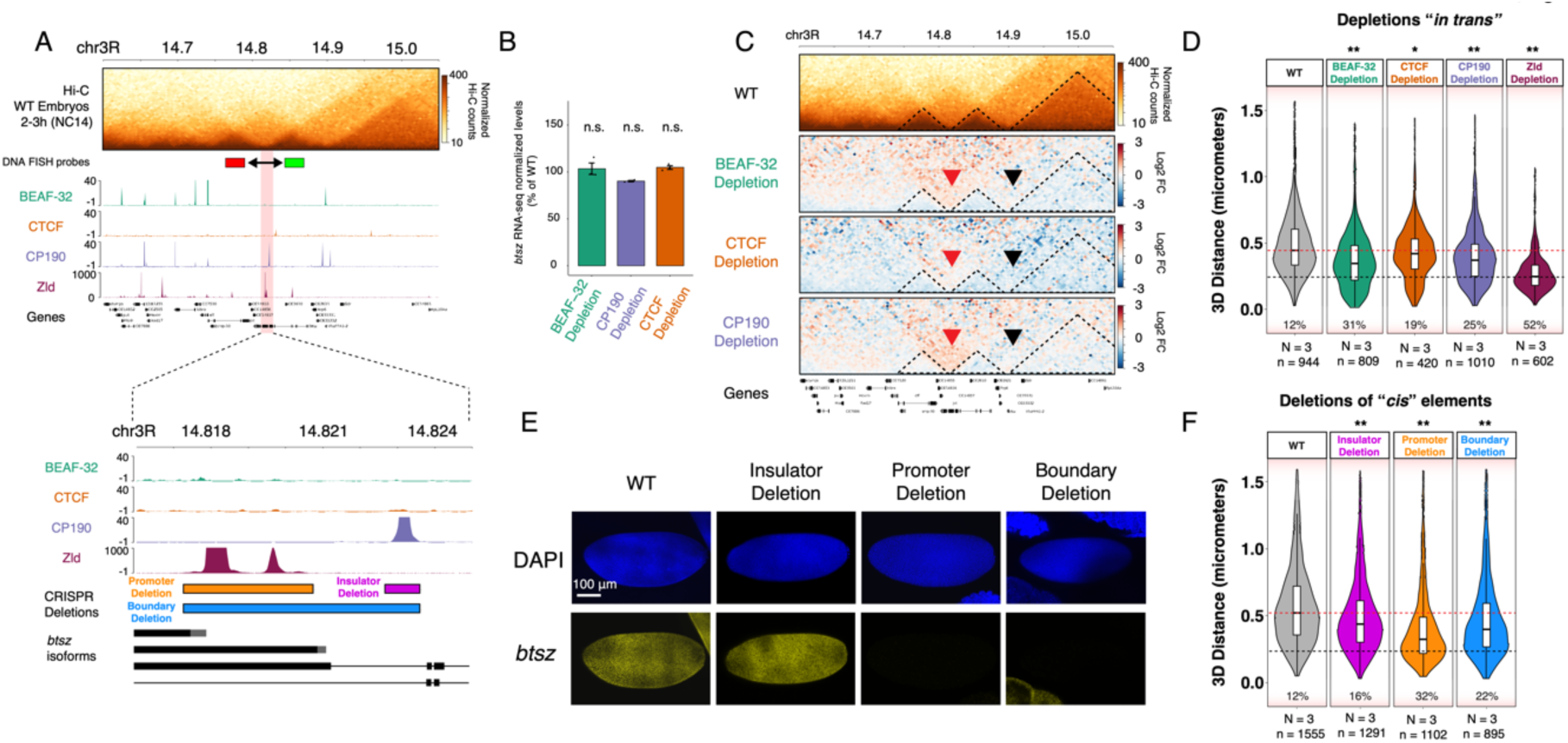
Active transcription is required for insulation at the *btsz* TAD boundary. **(A)** Genomic features at the *bitesize* (*btsz*) locus. *Top*: Hi-C matrix from WT embryos (2-3h), and BEAF-32, CTCF, CP190 occupancy (ChIP-seq from NC14 embryos) and Zelda occupancy (ChIP-seq from (*79*)). Location of DNA-FISH probes indicated by red and green rectangles. *Bottom*: Zoom in to occupancy at the TAD boundary and *btsz* promoter. Location of DNA regions deletions are indicated by colored rectangles. **(B)** Quantification of *btsz* RNA-seq signal in each insulator- depletion (normalized to WT); non significant (n.s.; FDR > 0.05). **(C)** Hi-C matrix from WT 2-3h embryos and differential Hi-C matrices (log2 Fold-Change) from 2-3h embryos (higher in insulator protein depletion: red, higher in WT: blue). Red arrow indicates the disrupted *btsz* boundary, black arrow the neighbouring unaffected boundary. **(D)** Quantification of 3D distances between the center of mass of the DNA-FISH probes indicated in (A) across multiple alleles from WT and insulator- depleted embryos at NC14 (N = number of embryos, n = number of alleles). Percentages indicate the number of alleles with distances between the two probes below 250 nm. Dotted lines indicates distances at the WT median (red) and <250nm (black),. Kolmogorov–Smirnov (KS) test, * p < 0.01, ** p < 0.001. **(E)** *In-situ* hybridization of *btsz* (yellow) in WT embryos and embryos with the CRISPR deletions indicated in (A) at NC14. DAPI shown in blue. Scale bar = 100 μm. **(F)** Quantification of 3D distances between the center of mass of the DNA-FISH probes indicated in (A) in WT and CRISPR deletion embryos. N = number of embryos, n = number of alleles measured. Percentages indicate the number of alleles with probe distances below 250 nm. Dotted lines indicate distances at the WT median (red) and 250nm (black). Kolmogorov–Smirnov (KS) test, * p < 0.01, ** p < 0.001.

However, there are examples of DEGs close to the disrupted TAD boundary, and therefore could be good candidates for enhancer hijacking – where an enhancer in one TAD leads to the mis-regulation of a gene in a neighbouring TAD after a disruption of the boundary that normally segregates the two. Such examples have been previously observed in both mammals (*9, 11, 66*) and *Drosophila* (*10, 67*). Here, using this more rapid depletion, in a single generation, we searched for enhancer hijacking events by looking for DEGs (zygotic-only) in the neighbouring TAD for TADs with affected boundaries after each depletion.

Taking the top 100 affected boundaries in each genotype, we identified 10 (3%; 10/325 DEGs), 34 (7.8%; 34/436) and 40 (6.7%; 40/597) cases in BEAF-32, CTCF and CP190 depletions, where a zygotic gene is upregulated in the neighbouring TAD (Table S4). We chose two of these potential hijacking candidates that also have characterised enhancers that are active at NC14 to explore in more detail: *wingless* and *short gastrulation* (Fig. 5D,E).

*Wingless* (*wg*) is a well-studied segment polarity gene whose expression is activated at NC14 to control anterior-posterior patterning of the early embryo. *wg* and its enhancers are contained between two CTCF/CP190 binding sites, which separate them from the neighbouring *wnt4* gene, which is only activated later in development (Fig. 5D). One of the CTCF/CP190 ChIP peaks overlaps a TAD boundary between *wg* and *Wnt4*. This peak is lost, and the TAD boundary disrupted after CP190 (and CTCF) depletion (Fig. 5D). Our RNA-seq experiments indicate no change in *wg* expression, but a slight yet significant increase in *wnt4* expression, located in the neighbouring TAD upon CP190 depletion (Fig. 5D). Using RNA fluorescent *in situ* hybridisation, we observed a few cells that acquire *wnt4* mis- expression, which largely co-localizes with the high levels of *wg* in a posterior stripe (Fig. 5F inset), or in a patch in the anterior of the embryo. This ectopic expression pattern suggests that the *wg* enhancer(s) controlling this expression pattern start(s) communicating with the *wnt4* promoter upon loss of CP190.

An even more dramatic example is at the *short gastrulation* (*sog)* locus. *sog* is an essential gene expressed in two bands along the anterior-posterior axis of the embryo that will give rise to the neuroectoderm, while been excluded from the presumptive mesoderm in between. Similarly to *wg*, the *sog* locus is flanked by CTCF and CP190 bound regions on both sides – the 3’ CTCF-CP190 bound region overlaps a disrupted domain boundary in both CTCF and CP190 depleted embryos (Fig. 5E). Our RNA-seq measurements showed no alteration in *sog* expression, but an up-regulation of two genes (*CG12708* and *CG15646*) in the neighbouring domain on the right hand-side in both CTCF and CP190 depleted embryos (Fig. 5E). *In situ* hybridisation using a probe that overlaps both genes shows that they are normally only expressed in a small domain in the anterior end of the embryo in NC14 stage embryos (Fig. 5G). However they become mis-expressed in a *sog*-like pattern in both CP190 and CTCF depleted embryos (Fig. 5G), again suggesting that a *sog* enhancer can now communicate with these genes’ promoters.

Taken together, our results indicate that these insulator proteins can regulate gene expression through multiple mechanisms at these stages of embryogenesis. This includes potentially acting directly at the promoter to activate gene expression (1-10% of DEGs) and by constraining enhancer activity at domain boundaries (3-8% of DEGs), exemplified by enhancer hijacking at the *wg* and *sog* loci. In these hijacking cases, neither the promoters (of both the patterning genes and mis-expressed genes (*wnt4*, *CG12708*, *CG15646*)), nor the enhancers (or their surrounding regions) are bound by insulators, indicating that these insulators do not mediate enhancer-promoter tethering, but rather modulate their communication indirectly, by maintaining the nearby boundary. However, both of these mechanisms can only account for a minority (<20%) of all DEGs. Interestingly, there are very few cases of DEGs that directly overlap (within 10kb) a disrupted TAD boundary, after insulator protein depletion. This indicates that TAD boundaries can be diminished while maintaining normal transcription of the housekeeping genes present at that boundary, and also perhaps that redundancy between insulator proteins and/or transcription itself helps to maintain insulation at boundaries.

### Transcription and insulator protein binding are both required for full insulation of a TAD boundary

To explore if gene transcription at a boundary can add to the overall robustness of the boundary, and thereby help maintain boundary insulation after depletion of insulator proteins, we functionally dissected one boundary. We selected a boundary that overlaps both an insulator bound region and an active promoter that drives strong expression of the main isoform of *bitesize* (*btsz*) during NC14 (*68*). This *btsz* TAD boundary (Fig. 6A) is occupied by the transcription factor Zelda (Zld) at NC14, which binds to the promoter of *btsz*’s main isoform, and by CP190, which binds ∼ 5 kb upstream (Fig. 6A). Both BEAF-32 and CTCF do not bind at the boundary, but CTCF binds to a site ∼12kb upstream of the promoter, while BEAF-32 co-binds with CP190 to the boundaries of the two adjacent domains (Fig. 6A).

Depletion of Zld leads to a strong downregulation of *btsz* expression (*22, 69*), which combined with Zld occupancy at the *btsz* promoter, strongly suggests that Zld directly activates the expression of this gene. In contrast, there is no significant change in *btsz* expression upon depletion of any of the three insulator proteins (Fig. 6B).

To measure the impact of insulator protein depletion on boundary function, we used both Hi-C and DNA-FISH. Differential Hi-C maps indicate an increase in interactions crossing the domain boundary upon depletion of any of the three insulator proteins (Fig. 6C). This was confirmed independently using Cromosight (diff_score in relation to WT: -0.61, -0.48 and -0.49 for BEAF-32, CTCF and CP190 depletions, respectively) and by calculating changes in insulation score (IS: - 0.71, -0.07, -0.04 and 0.13 for WT, BEAF-32, CTCF and CP190 depletions, respectively). Concordantly, DNA-FISH measurements of the distance between the centre of the two adjacent domains showed a decrease in domain distances (i.e. higher compaction) following insulator protein depletion (Fig. 6D): 12% of cells had these domains within 250nm in WT embryos, this shifted to 31%, 19% and 25% after the depletion of BEAF-32, CTCF and CP190, respectively. Thus, both techniques indicate that removing any of the three insulators impacts *btsz* domain boundary function, even if these insulators (BEAF-32 and CTCF) are not directly binding at the central domain boundary. A recent study proposed that CP190 acts predominately to insulate the subset (∼20%) of *Drosophila* boundaries that do not contain an active promoter (*29*). At least in the context of ZGA, our data indicates that CP190 can influence insulation at a TAD boundary containing an active promoter, although it is not the only requirement. Zld depletion led to an even stronger loss of boundary function (Fig. 6D, 52% of cells with distances <250nm), indicating that at this locus Zld binding and/or Zld regulated transcription is more important for boundary function than insulator binding.

To confirm that these effects are due to regulation in *cis*, and not to potentially secondary *trans* effects, we used CRISPR-Cas9 to delete three elements within the region: (1) the *btsz* promoter and promoter proximal region (∼ 3.5 kb), (2) the CP190 bound region (∼ 1 kb) leaving the *btsz* promoter intact and (3) both the *btsz* promoter and CP190 bound region (a 6 kb deletion spanning the entire boundary) (Fig. 6A, lower). To determine whether the deletions affected expression of *btsz*, we performed *in-situ* hybridisation. Deletion of the promoter (∼3.5kb) or the entire boundary (∼6kb) completely abolished *btsz* expression in NC14 embryos, as expected, while the insulator deletion had no detectable effect on *btsz* expression (Fig. 6E). Accordingly, since *btsz* is an essential gene, the promoter and the entire boundary deletions are homozygous lethal, while flies with the insulator binding site deletion are homozygous viable and fertile. To measure the insulation across the domain boundary in these lines, we performed DNA-FISH on embryos at NC14 (similar to Fig. 6D). Deletion of the insulator binding site led to a reduction in insulation between the two domains as seen by their increased proximity: the number of cells with distances <250 nm changed from 12% to 16% in the WT and insulator deletion, respectively (Fig. 6F). However, the changes were much more dramatic in embryos with the promoter deletion (Fig. 6F), changing form 12% (WT) to 32% (promoter deletion) of cells with distances <250nm. Therefore at this locus, deletion of the Zelda regulated promoter had a more pronounced effect on boundary function compared to deletion of the insulator bound region itself (Fig. 6F).

In summary, although deletion of the insulator binding site in *cis* (Fig. 6F), and depletion of insulator proteins (Fig. 6D) have an effect – the deletion of the promoter region and depletion of Zelda protein, which is essential for the activation of this promoter, had the most dramatic effect on boundary insulation. This suggests that transcription (or the occupancy of this promoter) is required for full boundary function, and acts together with insulator protein binding, perhaps to reinforce boundary insulation. Interestingly, of the insulators tested, the depletion of BEAF-32 had the most prominent effect (Fig. 6D) even though it is not bound to the central boundary. This indicates that insulator proteins can influence domain boundary function in a long-range manner at some loci, perhaps by perturbing the compaction of the neighbouring domains.

## DISCUSSION

### Dynamics of TAD establishment in early embryos

Previous studies using Hi-C in staged embryo pools showed that *Drosophila* TADs are first detectable during the major ZGA at NC14 (*22, 41*). Here, using an orthogonal single cell approach, DNA-FISH, in individual staged embryos, we confirm and extend these findings (Fig. 1). Although TADs are established during NC14, they are fuzzy due to extensive cell-to-cell heterogeneity and lower insulation compared to later developmental stages (Fig. 1, Supplementary Fig. S1). Our FISH data also shows that TADs are formed with similar temporal dynamics and cell-to-cell variability for domains with distinct transcriptional profiles, containing genes that are either inactive during ZGA (neutral TAD) or active (containing co-regulated or actively expressed genes that are not co-regulated) (Fig. 1). Therefore, although TAD formation coincides with the major wave of transcription during ZGA, transcription within the TAD does not seem to be required for the establishment of the TAD, in line with previous studies that blocked transcription using pharmacological inhibitors (*22, 70*). Conversely, transcription may be required to achieve full insulation at TAD boundaries containing a promoter, at least in the context of the *btsz* containing TAD (Fig. 6 and discussed below).

### TAD boundaries are a collection of diverse genomic elements in *Drosophila*

In mammals, deletions of individual CTCF sites typically has limited effects on TAD insulation. The fusion between two TADs usually requires deletion of multiple CTCF sites (*12, 51, 71*). In *Drosophila*, we observed, in line with others, that TAD boundaries are often occupied combinatorically by multiple insulator proteins, which can be a mixture between multiple peaks for the same factor and multiple peaks each for a different factor: e.g for NC14 see Fig. 2, and for later embryonic stages see (*4, 24–27*). For example, we observe both insulator co-binding at a single site (i.e. within a 200 bp window – Fig 2C, left) – consistent with the recruitment of CP190 by either BEAF-32 or CTCF (*32, 44, 49*), in addition to more distributed binding within the same boundary region (Fig. 2C, right), with up to 4 different peaks for the same factor within a 10kb boundary region. This suggests a potential basis for the robustness of TAD boundaries – perhaps the loss of a single factor is not enough to completely abolish boundary function. Notwithstanding, our data indicates that the strength of insulation is not simply dependent on the number of different co-bound insulator proteins. Different combinations of factors are associated with boundaries of different insulation strengths, with BEAF-32 appearing to act more redundantly, while CTCF & CP190 may act more cooperatively (Fig. 4F). These results highlight the diversity of *Drosophila* boundaries and the difficulty to predict whether an insulator-bound site will be able to form a boundary or not.

### Insulator proteins are individually not required for the formation of most TADs

Our depletion strategy allowed us to achieve near full depletion of these insulator proteins from chromatin, so they are not present at the moment when TADs are being established (Fig. 3). Using CUT&Tag with spike-ins was crucial to confidently and quantitatively compare insulator protein binding between control and insulator-depleted embryos. This revealed that there are no statistically significant peaks identified, of the thousands of wild-type peaks, in insulator depleted embryos. This is important, as even low levels of insulator protein is enough to sustain chromatin topology in other models (*17, 56*).

In the absence of the insulator proteins BEAF-32, CTCF or CP190, we observed that most *Drosophila* TADs can still form (Fig. 4), even though 86% of TAD boundaries are occupied by at least one of those proteins (Fig. 2). This robustness may be explained by alternative, not necessarily exclusive, hypotheses. One mechanism could be redundancy between insulator proteins, as multiple different proteins have been identified in *Drosophila*, as reviewed in (*23*). In mammals, TADs are formed by cohesin mediated loop extrusion, which stalls at CTCF bound sites (*15, 16*). Although there is currently no experimental evidence for cohesin mediated loop extrusion in *Drosophila*, cohesin could in principle be stalled by CTCF or any combination of insulator proteins in a similar manner. As we removed these three insulator proteins individually, other factors may compensate for their loss and facilitate stalling of loop extrusion to create TAD boundaries. Our data suggests that this may be the case for BEAF-32 – removal of BEAF-32 had minimal impact on boundaries co-bound by other insulator proteins, compared to boundaries occupied by BEAF-32 alone (Fig. 4F). However, this does not appear to be the case with CTCF and CP190 – depletion of either factor impacted boundaries that are occupied by many insulator proteins (3 or 4 factors), suggesting more cooperative interactions (Fig. 4F).

Alternatively, as the majority (almost 80% according to Ramirez et al., 2018) of *Drosophila* TAD boundaries overlap with active promoters of constitutively expressed genes (*4, 24, 35*), transcription may be involved in maintaining TAD boundary function. Yet two studies that used pharmacological inhibitors to block transcription did not observe effects on TADs or TAD boundaries (*22, 70*). On the other hand, our data shows that TAD insulation increases at later embryonic stages (Fig. 1E,G,I), which correlates with when many genes are expressed. Functional dissection of one boundary at the *btsz* TAD indicates that an active promoter is required for full boundary insulation, and had a stronger impact than the insulator bound region itself. This is suggestive of a role for active transcription in boundary insulation, although we cannot exclude that the functional ‘boundary entity’ is rather Pol II occupancy at the promoter (or formation of a pre-initiation complex) or Zelda binding itself. This promoter-associated function is also not the only requirement for boundary function (i.e. it does abrogate compaction completely), but perhaps it plays a role in the reinforcement of TAD boundaries after they are formed, which is consistent with the increase in insulation we observe as development proceeds.

Lastly, TAD boundaries may be strongly locus specific in *Drosophila*, i.e. each boundary may rely on a specific combination of factors in addition to the surrounding genomic context to function. This is supported by the different sub-sets of boundaries that are more sensitive to depletion of different factors, as we observed here during ZGA and in recent studies at different stages (*24, 29, 30, 39, 40, 72*). For example, for the factors depleted here, most disrupted boundaries were genotype-specific. Nevertheless, boundaries affected by depletion of CTCF and CP190 had more overlap compared to BEAF-32 depletion (Fig. 4). Even for the most affected TAD boundaries, the absence of the insulator protein did not lead to a complete loss of the TAD or complete fusion between neighboring TADs, but rather a weakening of the boundary. As TADs are just forming at this stage, and still quite fuzzy, we assumed that they would be easier to disrupt or break – but this does not seem to be the case. They are seemingly as resilient to perturbation here as seen in other contexts (*12, 51*).

### Insulator proteins contribute to gene expression through different mechanisms

By definition, the major event that is occurring during ZGA is the activation of transcription in the embryo, making this a very interesting stage to examine the requirement of these proteins for both chromatin topology and the initiation of gene expression. After the depletion of each of the three factors, a few hundred zygotically expressed genes had significant changes in their expression (Fig. 5). However, it’s interesting to note that this did not include many of the ‘classic’ minor and major wave early patterning genes (*58*). However, there are some exceptions: (1) 1 out of 10 pair-rule genes (*even-skipped (eve)* was slightly down-regulated in CP190 depletion, (2) 2 out of 8 homeotic genes had change expression: *Scr* was slightly up-regulated in BEAF-32 and down-regulated CTCF depletions, while *Ultrabithorax* (*Ubx*) was down-regulated in CP190 depletion (Table S3). None of the 13 gap genes were mis-regulated in any genotype. Therefore, ZGA can still occur largely unperturbed after the depletion of each of these insulator proteins, despite their occupancy at the majority of TAD boundaries and thousands of intra-TAD sites.

These insulator proteins appear to regulate a small fraction of genes (from 1-10% of the downregulated genes in each genotype) by directly binding at their promoter, which can be due to regulation of transcriptional initiation or mediating the communication with other cis-regulatory elements. We also identified a small number of potential cases of enhancer hijacking (3-8% depending on the genotype), where an enhancer in one TAD with a weakened boundary mis-appropriately activates a gene in the neighboring TAD (Fig. 5). For example, in the *wg* and *sog* loci, the TAD boundary seems to have restricted the enhancer’s activity such that weakening the boundary leads to enhancer activation of an additional target gene in the neighboring TAD. It is interesting to note that such enhancer hijacking cases varied in their effect size. In the *wg* locus, for example, disruption of the boundary is associated with a gain of *wnt4* expression in a very small number of cells. In comparison, disruption of a boundary at the *sog* locus led to *CG12708*/*CG15646* misexpression in the majority of *sog* expressing cells. These observations, together with recent studies indicate that the impact of boundary perturbations on gene expression, if any, depends on multiple factors (*73*). However, together both mechanisms (direct promoter regulation and enhancer hijacking) can only account for a minority (<20%) of the mis-expressed genes after these insulators depletion. DEG are generally not enriched near disrupted TAD boundaries - the median distance of mis-expressed genes is ∼125, 100 and 70 kb from a disrupted boundary for BEAF-32, CTCF and CP190 depletions, respectively (Fig. 5C). This suggests that these genes change in expression by other, as yet not understood, mechanisms.

As many insulator proteins, including CTCF in mammals, have been proposed to also act as ‘normal’ transcriptional activators, distinguishing between a role in *trans* versus a causal relationship in *cis* is difficult. Here, we used genetic deletion of regulatory element to dissect one complex TAD boundary that simultaneously overlaps an active promoter and an insulator bound region, during ZGA (Fig. 6). In that boundary, deleting the active promoter caused a stronger effect on TAD boundary insulation, in comparison to deleting the insulator binding site, which still had an effect on boundary insulation. We propose that TAD boundaries rely on distinct genetic features to achieve their maximum level of insulation. While at some loci, an active promoter (and/or transcription) may be the dominant factor for boundary function, at others insulator proteins may be more important. The interplay of these factors with the local chromatin environment is also likely to play a role in chromatin structure.

Taken together, our results suggest that throughout the *Drosophila* genome, the establishment of TADs is shaped by interactions at different levels, with different combinations of insulator proteins being required in a seemingly locus-specific manner and influencing the topology of each locus to a different degree. We hypothesize that redundant mechanisms, such as binding of multiple insulator proteins, coupled with transcriptional activity (or active promoters), underly the robustness of genome structure. Future studies are needed to shed light on the relationship between insulator combinatorial binding and their genomic context to dissect the rules that governs the formation of domain boundaries in *Drosophila*, and the cross-talk between boundary function and transcription.

## METHODS

The western blots, fluorescent *in-situ* hybridisations, Hi-C, RNA-seq, ChIP-seq data processing, RNA-seq analysis, and Hi-C data analysis were performed with standard procedures. Detailed Methods for each are provided in the Supplementary Methods, while the more specific methods used in this study are provided below.

### Genetic deletion/knockdown of CTCF, BEAF-32 and CP190

Maternal knockout of CTCF depletion was done as described previously in (*39*). Briefly, CTCF knockout flies were rescued with an FRT-flanked 5 kb CTCF genomic rescue transgene and developed into viable and fertile adults. The excision of the CTCF rescue cassette from male and female germlines was achieved through nanos-GAL4:VP16 (NGVP16)-driven expression of UAS-FLP, and the resulting maternal/zygotic CTCF-depleted embryos were collected. This thereby generates complete genetic loss-of-function embryos.

BEAF-32 and CP190 depletion was done by RNAi-mediated knockdown using stocks carrying shRNAs as described previously (*55*). We first compared the efficiency of different germline GAL4 drivers and different temperatures. Virgin females carrying either BEAF-32 or CP190 shRNA (VDRC #330274 and BDSC #33903, respectively) were crossed to males from either the MTD-Gal4 (BDSC #31777) or Mat-tub-Gal4 lines (BDSC #7063). These GAL4 lines express GAL4 either during all stages of oogenesis (MTD-Gal4) or only during late stages (Mat-tub-Gal4) (*55*). The crosses were incubated at 25°C or 29°C, leading to different GAL4 efficiencies. F1 virgin females carrying one copy of either the BEAF-32 or CP190 shRNAs and one copy of the GAL4 transgene(s) were crossed to males carrying the BEAF-32 or CP190 shRNA respectively. F2 embryos were collected and used for all experiments. For BEAF-32 depletion, the optimal combination was the mat-tub-GAL4 driver and 25°C, as other conditions (either the MTD- gal4 driver or the 29°C temperature) highly increased sterility. For CP190 depletions, the MTD-Gal4 driver and 29°C was used as this resulted in stronger levels of protein depletion (assessed by western blot).

### CRISPR deletions in the *btsz* locus

To generate flies with CRISPR deletions (Fig. 6), CRISPR donor and gRNA plasmids were constructed following the strategy for “gene replacement with pHD-DsRed-attP” described in (*74*). gRNA sequences were generated by annealed oligo cloning and inserted into the BbsI site of the pU6- BbsI-gRNA vector. To generate the homology arms, we PCR-amplified regions between 2-3kb starting directly from the upstream or downstream cutting site of each gRNA, and inserted those into the SapI and AarI sites of the pHD-DsRed- attP vector, using the In-Fusion cloning kit (#639650, Takara Bio USA, Inc.). Primers and gRNA oligos are listed in Table S5. Both gRNA and homology repair template plasmids were injected into embryos carrying a *vasa-Cas9* transgene on the 3^rd^ chromosome (BDSC # 51324, w[1118]; PBac{y[+mDint2]=vas-Cas9}). Hatching adults were crossed to flies carrying a 3^rd^ chromosome balancer marked by GFP, and the progeny was screened for both the DsRed and GFP markers. PCR-genotyping was used to confirm the editing. DsRed+/GFP+ siblings were crossed, and the progeny was assessed for viability. Flies carrying a disruption of the *btsz* promoter or the whole boundary were not able to be maintained as homozygous stocks and were therefore maintained as a trans-heterozygous stock over a marker balancer chromosome.

### Embryo collections

Freshly hatched adults were placed in embryo collection vials with standard apple cap plates. For DNA-FISH experiments, following three 1 h pre-lays, the flies were allowed to lay for 3h and embryos were directly collected. For genomics experiments, following three 1 h pre-lays, the flies were either allowed to lay for 30 min after which the embryos were aged for 2h10 to reach the interval 2h10-2h40h (ChIP-seq at NC14), or the flies were allowed to lay for 1 h, after which the embryos were aged for 2 hours to reach the interval 2-3h (CUT&Tag, Hi-C, Western Blot and RNA-seq). The embryos were then dechorionated using 50% bleach, and washed with deionized water (dH2O) and PBT 0.1% (phosphate buffered saline (PBS) containing 0.1% Triton X-100). Embryos used for FISH were cross-linked in 4% formaldehyde for 20 min at room temperature, de-vitelinized and stored in 100% methanol at -20°C. Embryos used for Western blot and RNA-Seq were kept on ice-cold PBT, and NC14 embryos were manually selected using an embryo needle, based on morphological indicators (*75*) under a stereoscope and then directly placed in sample buffer (Western Blot) or snap-frozen in liquid nitrogen (RNA-seq). Embryos used for ChIP-seq, CUT&Tag and Hi-C (2-3hr) were crosslinked in 1.8% formaldehyde for 15 min or 3% formaldehyde for 30 min at room temperature. Fixation was stopped by addition of PBT 0.1% + Glycine 125mM, followed by a wash with PBT 0.1%, then air-dried on tissue and snap-frozen in liquid nitrogen.

### 3D DNA FISH and combined DNA/RNA FISH

DNA FISH probes were generated by PCR amplification from *D. melanogaster* genomic DNA (7 – 8.5 kb fragments) and TA cloning into a PGEMT-Easy vector (Promega #A1360) (primers listed in Table S5), except for the *btsz* locus, for which BACs were used from DGRC (CHORI22-94I16 and CHORI22-115C08). The Nick Translation kit (Abbott Bioscience #7J0001) was used to fluorescently-label probes.

Embryos fixed with formaldehyde 1.8% and stored in 100% methanol were re-hydrated, washed three times in 2x SSCT (2x Saline Sodium Citrate buffer (SSC) + 0.1% Tween), and then washed once in 20% formamide and in 50% formamide (both in 2X SSCT), at room temperature on a rotating shaker. This was followed by two one-hour washes in 50% formamide at 37°C while rotating. The 50% formamide was removed, and the embryos were denatured at 80°C for 15 minutes in a water bath, placed on ice and mixed with hybridization mix containing the fluorescent DNA probes. Following overnight hybridization at 37°C, embryos were washed 2x in 50% formamide while rotating at 37°C, 1x with 20% formamide and 3x in SSCT while rotating at room temperature. For DNA FISH using a fluorescent marker (Fig. 6, *btsz* CRISPR deletions), the protocol continued using the HCR RNA FISH (Molecular Instruments) protocol to detect the GFP balancer chromosome, following manufacturer’s instructions. Embryos were placed in ProLong™ Gold mounting medium with DAPI (ThermoFisher Scientific #P36931), and mounted onto a slide.

Slides were imaged using a Leica SP8 confocal microscope with a 100x objective (HC PL APO CS2 100x / NA 1.4 / Oil), a 405 nm laser, a white-light laser (470-670 nm), and HyD detectors. The z-stack step size was 200 nm. For all DNA FISH samples we acquired z-stacks covering a single layer of nuclei in the center of NC14 embryos. At least three embryos, and hundreds of alleles, were used per condition.

To precisely stage each single embryo, the number of nuclei in a 50 μm^2^ window was counted in each image, according to (*76*). Images were deconvolved using the Huygens Professional software (SVI) with default parameters. For the quantification of distances between DNA FISH probes in each image, we used a custom FIJI plugin (“Analyze FISH spots”) developed in house by EMBL’s Advanced Light Microscopy Facility (ALMF). The plugin has two main functions: (1) detect spots across different channels and (2) automatically calculate the 3D distances between spots. Briefly, the x, y and z coordinates of FISH spots are determined based on a manually provided value for signal intensity and background in each channel following visual inspection. The plugin displays the spots in the image and the multi-point tool is used to manually select “clusters” of nuclear spots within a nucleus in the different channels along the z-stack (two or three channels depending on the number of DNA FISH probes used). After all spot clusters are manually selected, the FIJI plugin calculates the pair-wise 3D distances between the Center-of-Mass (CoM) of each spot/channel in all clusters. These distances were used in the DNA FISH violin plots throughout this study. A Kolmogorov-Smirnov test was used to compare the distribution of distances between a given genotype and WT samples.

### ChIP-seq and CUT&Tag of insulator proteins in NC14 embryos

ChIP-seq was performed as described in (*77*). After sonication and chromatin extraction, the chromatin was aliquoted into fresh tubes and stored at -80°C until use. The quality of the sheared chromatin was determined by agarose gel electrophoresis to observe chromatin fragment size distribution. The following antibodies were used; rabbit anti-CTCF (gift from R. Reinkawitz), rat anti-CP190 (gift from P. Georgiev), goat anti-Su(Hw) (gift from P. Geyer) and rabbit anti-GAF (gift from J. Lis), which were incubated overnight with chromatin in RIPA buffer (140mM NaCl, 10 mM Tris-HCl pH 8.0, 1 mM EDTA, 1% Triton X-100, 0.1% SDS, 0.1% Na-deoxycholate, 1x Roche cOmplete Protease inhibitors) in a total volume of 900 μl. We used 6 ug of chromatin for CTCF, 4 ug for CP190, 2 ug for Su(Hw) and 10 ug for GAF. Chromatin was fixed with 1.8% formaldehyde for 15 min for CTCF and Su(Hw) and with 3% formaldehyde for 30 min for CP190 and GAF. The next day 25 μl of magnetic protein A/G beads (Dynabeads, Invitrogen, 10002D and 10004D) were washed with 1ml of RIPA buffer and added to the IPs for an additional 3 hour incubation on the rotating wheel at 4°C. For the BEAF-32 ChIP, 25 μl of protein G beads were combined with 100ul of the BEAF-32 antibody (DSHB, #1553420) and 300 μl RIPA buffer for 2 hrs. This was followed by two washes with RIPA and resuspension in 100 μl of RIPA, which was added to the purified chromatin and incubated on the rotating wheel at 4°C overnight. The ChIPs were then washed for 10 min on the rotating wheel with 1x 1 ml RIPA, 4x 1 ml RIPA- 500 (500 mM NaCl, 10 mM Tris-HCl pH 8.0, 1 mM EDTA, 1% Triton X-100, 0.1% SDS, 0.1% Na-deoxycholate, 1x Roche cOmplete Protease inhibitors), 1x 1 ml LiCL buffer (250 mM LiCl, 10 mM Tris-HCl pH 8.0, 1 mM EDTA, 0.5% IGEPAL CA-630 CA-630, 0.5 % Na-deoxycholate) and 2x 1 ml TE buffer (10 mM Tris pH 8.0, 1 mM EDTA) on a magnetic rack in the cold room. The chromatin was then RNase-treated (#10109142001, Roche), reverse cross-linked overnight with 0.5 mg/ml Proteinase K and 0.5% SDS at 65°C. The next day the DNA was purified with Phenol-Chloroform purification and precipitated with ethanol, Sodium Acetate pH 5.3 and glycogen to obtain pure DNA. Library preparation was performed using the NEBNext® Ultra™ II DNA Library Prep Kit for Illumina® (NEB #E7645S), following the manufacturer’s instructions.

For quantitative CUT&Tag spike-in with the same number of *D. virilis* nuclei was used. Nuclei were counted using the BD LSRFortessaTM X-20 Flow Cytometer at the EMBL Flow Cytometry Facility and snap-frozen. We thawed fixed *Drosophila melanogaster* and *Drosophila virilis* nuclei (50,000 each) and then mixed the nuclei from both species for a total of 100,000 nuclei/sample. A conA bead slurry was added to the sample containing both *D. melanogaster* and *D. virilis* nuclei and placed on a rotating wheel for 10 min at 4°C. The nuclei-beads complex was washed, permeabilized and 1 μl of primary antibody was added to each sample. For the CTCF and CP190 CUT&Tag experiments we used the same antibodies as listed for ChIP-seq experiments, and for the BEAF-32 CUT&Tag we used a primary antibody gifted by C. Hart. The tubes were placed on a rotating wheel and slowly rotated (5 rpm) overnight at 4 °C. A secondary antibody solution (1:100) was added to the beads and incubated on a nutator at room temperature for 1h. We used the following secondary antibodies: Guinea Pig anti-Rabbit IgG (H+L) (Antibodies-online, ABIN6923140), Rabbit anti-Rat IgG (H+L) (Thermo Fisher Scientific, A18917) and Rabbit Anti-Mouse IgG (H+L) (Abcam, ab46540).

Samples were then washed 3x and the CUT&Tag was performed as described previously (*78*). The PCR products were purified with Agencourt AMPure XP beads (Beckman Coulter, #A63881), and quantified with Qubit and ran on the Bioanalyzer using hs DNA reagents and chips. Final libraries were multiplexed and sequenced with 75 bp paired-end reads using a Illumina NextSeq 500 platform at the EMBL Genomics Core Facility.

### Identifying disrupted TAD boundaries using Chromosight and Pareidolia

TAD boundary changes in 10 Kb bin cooler matrices between wild-type and insulator- depleted samples were detected using Chromosight (*57*) and quantified with the pareidolia tool (https://github.com/koszullab/pareidolia) (subsample=True, density_thresh=None, pearson_thresh=0.0, cnr_thresh=0.0) using the 10Kb TAD boundaries as defined by HiCExplorer. Pareidolia quantifies the correlation of the Hi-C signal to the expected kernel (here for TAD boundaries) and report the difference (depletion minus wild-type, producing a negative score upon loss of correlation) together with a signal-to-noise ratio (snr) score indicated how good was the separation between signal and noise in the evaluated sub-matrix. TAD boundaries with a snr less than 5 were excluded. As pareidolia does not provide any significance value on the reported differential score we opted for a stratification approach where we compared the top changing boundaries (here 100) to a set of 200 “stable” boundaries selected as the 200 boundaries with the smallest absolute differential score.

## ACKNOWLEDGEMENTS

The authors thank all Furlong lab members for very helpful discussions and comments. This work was technically supported by the EMBL’s Genomics core facility and by the public FlyBase and RedFly databases. We are very grateful to the following groups for kindly providing antibodies: Rainer Renkawitz (CTCF), Pavel Georgiev (CP190), Pamela Geyer (Su(Hw)), John Lis (GAF), Craig Hart (BEAF-32) and Chris Rushlow for kindly providing Zelda mutant flies.

## Author Contributions

G.R.C, R.R.V, T.P and S.F. performed experiments. C.G. performed computational analyses, including ChIP-seq, CUT&Tag, Hi-C analyses and integration between the datasets. T.B.N.C and P. L. performed Hi-C (Chromosight) analyses. A.R. performed RNA-seq analyses. S.F. generated CRISPR deletion lines. G.R.C and E.E.F. conceptualised the project, designed and analyzed experiments, and wrote the manuscript with input from the authors.

## Data availability

All sequencing data (Hi-C, ChIP-seq, CUT&Tag, RNA-seq) from this study were deposited in EBI ArrayExpress with accession numbers E-MTAB-9158 (Hi-C), E-MTAB-9156 (ChIP-seq), E-MTAB-11993 (CUT&Tag) and E-MTAB-11978 (RNA-seq).

